# DnaK duplicates and regionally evolves for the increase of proteomic complexity in bacteria

**DOI:** 10.1101/2023.06.19.545549

**Authors:** Zhuo Pan, Li Zhuo, Tian-yu Wan, Rui-yun Chen, Yue-zhong Li

## Abstract

Hsp70 is important for organismic cells to maintain proteostasis and the chaperone protein is duplicated in all eukaryotes and many prokaryotes. Although the functioning mechanism of Hsp70 has been clearly illuminated, the chaperone duplication and functional evolution has been less investigated. DnaK is a highly conserved bacterial Hsp70 family. Here we showed that the *dnaK* gene is present in 98.9% bacteria and 6.4% bacteria possess duplicated *dnaK*s; the occurrence and duplication is positively corelated to the increase of proteomic complexity. We identified the interactomes of the two DnaK paralogs in *Myxococcus xanthus* DK1622, which are mostly nonoverlapped, but both preferring the α&β domain proteins. Consistent with the proteomes, the MxDnaK substrates are both significantly size-larger and pI-higher than that of the single *E. coli* DnaK. MxDnaK1 is heat-shock inducible, prefers to bind cytosolic proteins, while MxDnaK2 is decreased by heat shock, and is more associated with membrane proteins. The nucleotide binding domain and the substrate binding beta domain are responsible for the significant changes of DnaK substrates, and the former also determines the dimerization of MxDnaK2, but not MxDnaK1. Our work highlights that DnaK is duplicated and regionally evolved for the increase of proteomic complexity in bacteria.

## Introduction

The 70-kDa heat shock protein (Hsp70) is a kind of ATP-dependent molecular chaperones. Hsp70s participate in a wide range of cellular processes by, in the cooperation with other chaperones, folding *de novo* proteins, refolding or degrading misfolded proteins produced under heat or oxidative stress ^1–3^. A typical Hsp70 protein consists of three functional regions: An N-terminal 45-kDa nucleotide binding domain (NBD), which is linked with a 15-kDa substrate binding domain (SBDβ) and then a 10-kDa α-helical lid domain (SBDα). Functionally, NBD binds with ATP, SBDβ interacts with peptide substrates, and SBDα functions as a lid to trap substrates in the polypeptide binding cavity. In addition, some Hsp70s have a highly disordered C-terminal tail (CTT), which directly precedes the lid domain with an unclear function ^4, 5^. Two types of cochaperones involve in the Hsp70 function: The J domain protein (JDP), which stimulates the ATPase activity to assist Hsp70 chaperone to fold substrates ^6^, and the nucleotide exchange factor (NEF), which promotes ADP dissociation and the release of folded substrates ^7^. JDP binds at the interface between the NBD and the SBDβ, while NEF binds to the NBD region to function ^8^. After releasing the folded proteins and ADP, Hsp70 binds new proteins and ATP to restart the allosteric cycle ^9^.

Based on the conserved domain, the Hsp70 proteins are divided into two superfamilies ^10^. The cd10170 covers the NBD domain only, while cl35085 stretches over the NBD and SBD regions. In the cl35085 protein superfamily, some proteins are highly conserved and are exclusively clustered, forming the PRK00290 family, and these proteins are usually characterized as canonical Hsp70 chaperones (DnaKs in bacteria). The Hsp70s have multiple copies in cells of all eukaryotes and ∼40% prokaryotes ^8, 10^. The Hsp70 duplicates usually play divergent cellular functions, but with an overlap to different extents depending on the similarity of the paralogs ^11–13^. For example, human HSC70 and HSP70 are 85% identical, and the phenotypes of their knockout mutants are different, which is consistent with their largely nonoverlapping substrates and highly distinct expression patterns ^14^. Ssb1 and Ssb2 are two Hsp70s of *Saccharomyces cerevisiae*, sharing an almost identical sequence and thus an almost identical nascent substrate pool ^15^; the deletion of *Ssb1* or *Ssb2* is phenotypically silent, but double deletion of the two genes causes slow growth and sensitivity against cold ^16^. Hsp70 also duplicates and plays distinct functions in many bacteria, including the pathophysiology of some serious pathogens ^17–19^. The functional diversification of distinct Hsp70 isoforms in cell is suggested to attribute to differential expression, abundance, subcellular localization and substrate specificity ^8^, which, however, has been less investigated. For example, in addition to the above human and yeast, only two other organisms, i.e. *Escherichia coli* and *Mycobacterium smegmatis* have reported of the DnaK clients, which are significant different not only in the substrate number but also in the composition ^20, 21^.

*Myxococcus xanthus* DK1622 is the model strain of myxobacteria. The Gram-negative bacterium possesses a large set of duplicate genes for the complex social behavior and environment adaptation ^22, 23^. Our previous studies showed that, in *M. xanthus* DK1622, nine genes encode the cl35085 proteins (including two PRK00290 proteins), and six encode the cd10170 proteins ^10^. The two PRK00290 proteins (MXAN_3192 and MXAN_6671) are phylogenetically located in the same subbranch with the typical DnaK proteins of other bacteria like *E. coli*, *Bacillus subtilis* and *Lactococcus lactis*, while the other cl35085 proteins and the cd10170 proteins are in different branches out of these typical DnaKs. The two PRK00290 proteins, not the other *Myxococcus* Hsp70s, could alternatively compensate the functions of EcDnaK (DnaK of *E. coli*) for growth ^10^. MXAN_3192 (MxDnaK1), the only undeletable *hsp70* gene in DK1622, is expressed at an extremely high level and is inducible by heat shock. In comparison, the transcription of MXAN_6671 is low and decreases in response to heat shock. MXAN_6671 is deletable, and the deletion makes the mutant sensitive to oxidative stress and deficient in some social behaviors ^10, 24, 25^. These results suggested that the two *Myxococcus* DnaKs are functionally divergent, but with an elusive mechanism.

In this study, we surveyed the distribution of DnaKs in prokaryotes and found that the DnaK chaperone is duplicated with the increase of cell proteomic complexity in bacteria. We identified the interactomes of the two DnaKs in *M. xanthus* DK1622, compared the two interactomes by themselves and with that of EcDnaK, performed region inter-swapping between MxDnaKs, and analyzed their interactome and cellular phenotype changes. The substrate spectrums of DnaKs possess highly similar SCOP (structural classification of proteins) folds, but the DnaK duplicates recognize largely nonoverlapping substrates. The functionality of DnaK duplicates is mainly associated with the regional evolution of NBD and SBDβ, which drive the functional specificity.

## Results

### DnaK duplicates for the increase of proteomic complexity in bacteria

We screened the existence of proteins with the PRK00290 domain in the genomes of representative prokaryotes and obtained a total of 16947 DnaKs, 16540 from 15554 bacterial genomes and 407 from 529 archaeal genomes (**Supplementary dataset 1**). The number of sequenced genomes varied greatly in different taxa, for example, the phyla of *Proteobacteria*, *Actinobacteria*, *Firmicutes* and *Bacteroidetes* occupied 89.4% of the total 16083 prokaryotic genomes. Distribution of the DnaKs also greatly varied (**Table S1**). Totally, 98.9% of the bacterial genomes contained at least one *dnaK* gene, while the *dnaK* gene was present in 75.6% of the archaeal genomes. Many archaeal genomes lacked *dnaK*, for example, there was no *dnaK* gene in the 58 *Crenarchaeota* genomes, but the *Euryarchaeota* genomes might possess none, single, or multiple *dnaK* genes. **Figure 1A** exhibits the phylogenetic tree of the 16947 DnaKs. The DnaK phylogeny was normally in accordance with the phylogeny of prokaryotic organisms, but with some exceptions. For example, the *Cyanobacteria* DnaK clade contained some DnaK branches of *Firmicutes*. The *Euryarcharota* DnaKs were divided into three groups (blue branches in **Figure 1A**), which were mixed with the *Firmicutes* DnaKs, suggesting their possible evolutionary relationships.

**Figure 1.**
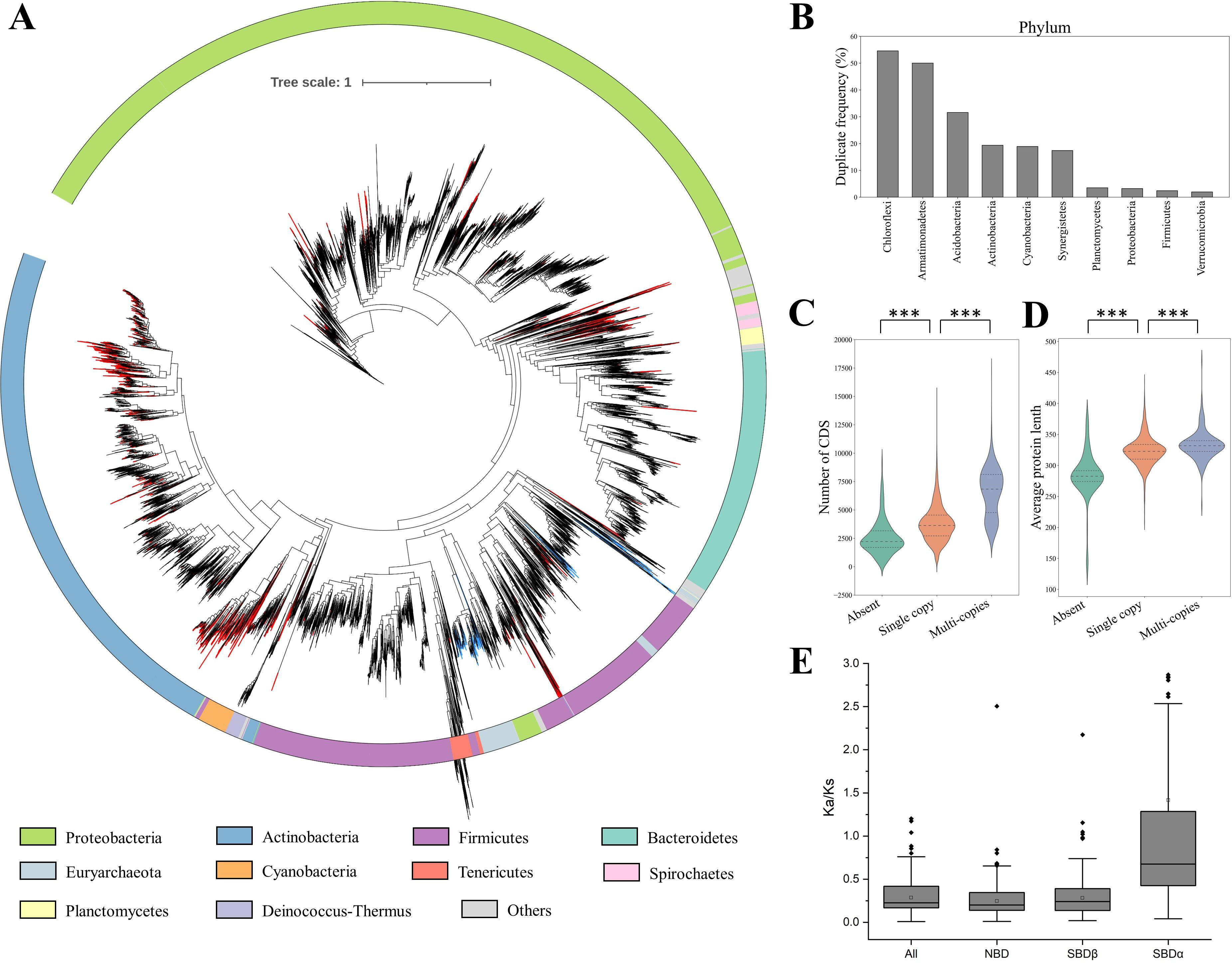
The distribution and characteristic analysis of prokaryotic DnaKs. (**A**) Phylogenetic analysis of the 16947 DnaK proteins from representative prokaryotes based on the neighbor-joining method by using the whole sequence of DnaK proteins. Multiple copies of DnaKs in a single genome are marked with red branches, and the *Eurarcharota* DnaKs are marked with blue branches. The top ten phylum resources of DnaK are colored in the outer ring, while the others are in grey. (**B**) The top 10 phyla with high ratios of duplicate *dnaK*s in prokaryotes. (**C**-**D**) The CDS number (**C**) and the average protein length (**D**) of genomes with none, single copy and multiple copies of *dnaK*. The p value is based on the Mann‒Whitney test. ***p < 0.001. (**E**) The Ka/Ks values of the whole protein sequences and the NBD, SBDβ and SBDα region sequences of all duplicate *dnaK* pairs.

There were 999 representative genomes of bacteria (6.4% of the total) possessing two or more *dnaK*s, and seven archaea encoding multiple DnaK proteins. Phylogenetic analysis showed that the distribution of DnaK duplicates was considerably wide but uneven in prokaryotes (red branches in **Figure 1A**). At the phylum level, five bacterial phyla had a high ratio with multiple copies of the *dnaK* gene (the phyla containing less than 10 genomes were not counted); *Chloroflexi* contained the highest *dnaK* duplication ratio (54.5% of 55 genomes), followed by *Acidobacteria* (31.6% of 38), *Actinobacteria* (19.4% in 3247), *Cyanobacteria* (18.9% in 180), and *Synergistetes* (17.4% of 23) (**Figure 1B**). The other phyla had a low ratio (< 3.5%) or none of duplicate *dnaK*s (**Table S1**). The duplication ratio of *dnaK* in some lower taxonomic units of bacteria might greatly increase, even reaching one hundred percent (**Figure S1**). For example, at the class level, the duplication ratios of *dnaK* in the *Ktedonobacteria* and *Chloroflexia* of the *Chloroflexi* phylum counted up to 88.2% and 72.7%, respectively. At the order level, the *dnaK* gene was duplicated in all the sequenced genomes of *Chloroflexales* (an order of *Chloroflexia*) or *Syntrophobacterales* of *Deltaproteobacteria*.

Notably, in the *dnaK*-free genomes, the median number of coding sequences (CDS) was 2220, and the upper and lower quartiles were 1698 and 3164, respectively (**Figure 1C**). In comparison, the median CDS number was 3629 (2715-4549) in the genomes with single copy of *dnaK*, and 6832 (4766-8123) in the genomes with multiple copies of *dnaK*. Obviously, the CDS number was significantly different between the genomes with none and single copy, or single copy and multiple copies of *dnaK* (p < 0.001). Similarly, in terms of protein size, the proteomes of prokaryotes with multiple copies of *dnaK* (332 (323-340)) were significantly larger than those with one copy of *dnaK* (323 (310-334)), which were further larger than those of the *dnaK*-free prokaryotes (283 (274-292)) (**Figure 1D**; p < 0.001). These results strongly suggested that the occurrence and duplication of *dnaK* are in accordance with the increase of proteomic complexity, thus helping host cells to deal with more and larger proteins.

In the prokaryotes with DnaK duplicate pairs (more than two DnaK duplicates in a genome were not analyzed), with a few exceptions, the Ka/Ks value of SBDα was almost always higher than that of NBD or SBDβ (**Supplementary dataset 2**). After removing the exceptions (collected in a separate table of **Supplementary dataset 2**), the median Ka/Ks value was 0.22 (0.17-0.42) for the whole protein sequence, 0.20 (0.14-0.35) for NBD, 0.24 (0.14-0.39) for SBDβ, and 0.68 (0.43-1.28) for SBDα (**Figure 1E**). Obviously, NBD was the most conserved region, while SBDα varied greatly. The duplicate DnaKs are thus regionally evolved mostly in a similar pattern.

Myxobacteria are a group of Gram-negative bacteria characterized by social behavior and complex life cycle. The myxobacteria have been recently upgraded from the *Myxococcales* order in *Deltaproteobacteria* to the *Myxococota* phylum based on 120 conserved single-copy marker genes as well as rRNA genes ^26^. Totally, 93 DnaKs were revealed in the 48 completely sequenced genomes of myxobacteria; 43 strains encoded two DnaKs, four strains, i.e. *Nannocystis exedens* DSM 71, *Corallococcus exiguous* NCCRE002, *Nannocystis pusilla* DSM 53165, and *Pajaroellobacter abortibovis* BTF92-0548A/99-0131, encoded a single DnaK, and one strain (*Vulgatibacter incomptus* DSM 27710) encoded three. The DnaK duplicates of myxobacteria were phylogenetically separately located in two clades, and the phylogeny of DnaKs in each clade was highly consistent with that of the phylogenomic tree (**Figure S2**). The missing DnaK in the four single DnaK-containing strains was the DnaK1 homologue of *M. xanthus* DK1622 (MxDnaK1), while the third DnaK in *V. incomptus* DSM 27710 was the MxDnaK2 homologue. The phylogenetic analysis indicated that the duplication of DnaK in myxobacteria occurred in the early days of evolution, and the functional importance of the DnaK duplicates might vary in different myxobacterial taxa.

### Interactomes of MxDnaK1 and MxDnaK2, compared with that of EcDnaK

To explore the functional evolution of DnaK duplicates, we identified and compared the interactomes of the two DnaKs of *Myxococcus xanthus* DK1622, as well as with that of *Escherichia coli* DnaK. MXAN_3192 and MXAN_6671 in *M. xanthus* DK1622 are the two Hsp70 proteins with the PRK00290 domain, named MxDnaK1 and MxDnaK2 in this study. The MxDnaK1 and MxDnaK2 proteins share 58% and 62% identities with the EcDnaK of their amino acid sequences, respectively. The two proteins themselves have a total of ∼60% identity of the amino acid sequences, and the NBD, SBDβ and SBDα regions share the identity of 64%, 70% and 30%, respectively (**Figure S3**). Thus, the SBDα region has the highest variation among the three regions of MxDnaK1 and MxDnaK2.

We performed coimmunoprecipitation (Co-IP) experiments to identify the interacting proteins of MxDnaK1 and MxDnaK2 (**Figure 2A**). The purified SBDα fragments of MxDnaK1 and MxDnaK2 were employed as the antigens to generate antibodies specific to the two DnaKs. According to the western blotting results with the soluble lysate of *M. xanthus* DK1622 cells, as well as a MxDnaK1 mutant swapped with the MxDnaK2 SBDα region (YL2204, see below), the two antibodies precisely recognized MxDnaK1 and MxDnaK2, respectively (**Figures 2B**, **S4** and **S5**). To stabilize the DnaK-substrate complexes, a high concentration of apyrase was added to deplete ATP in the lysis buffer, allowing the Hsp70 cycle to be hampered at the ADP-bound state ^27^. Negative controls for each MxDnaK (see methods) were employed to exclude the non-specifically bound proteins of beads and antibodies. Consequently, a total of 546 MxDnaK1 interactors and 375 MxDnaK2 interactors, including the cochaperones JDPs and NEFs, were identified by LC‒MS/MS with q value ≤ 0.01. Interestingly, the two interactomes were largely nonoverlapped (**Supplementary dataset 3**).

**Figure 2.**
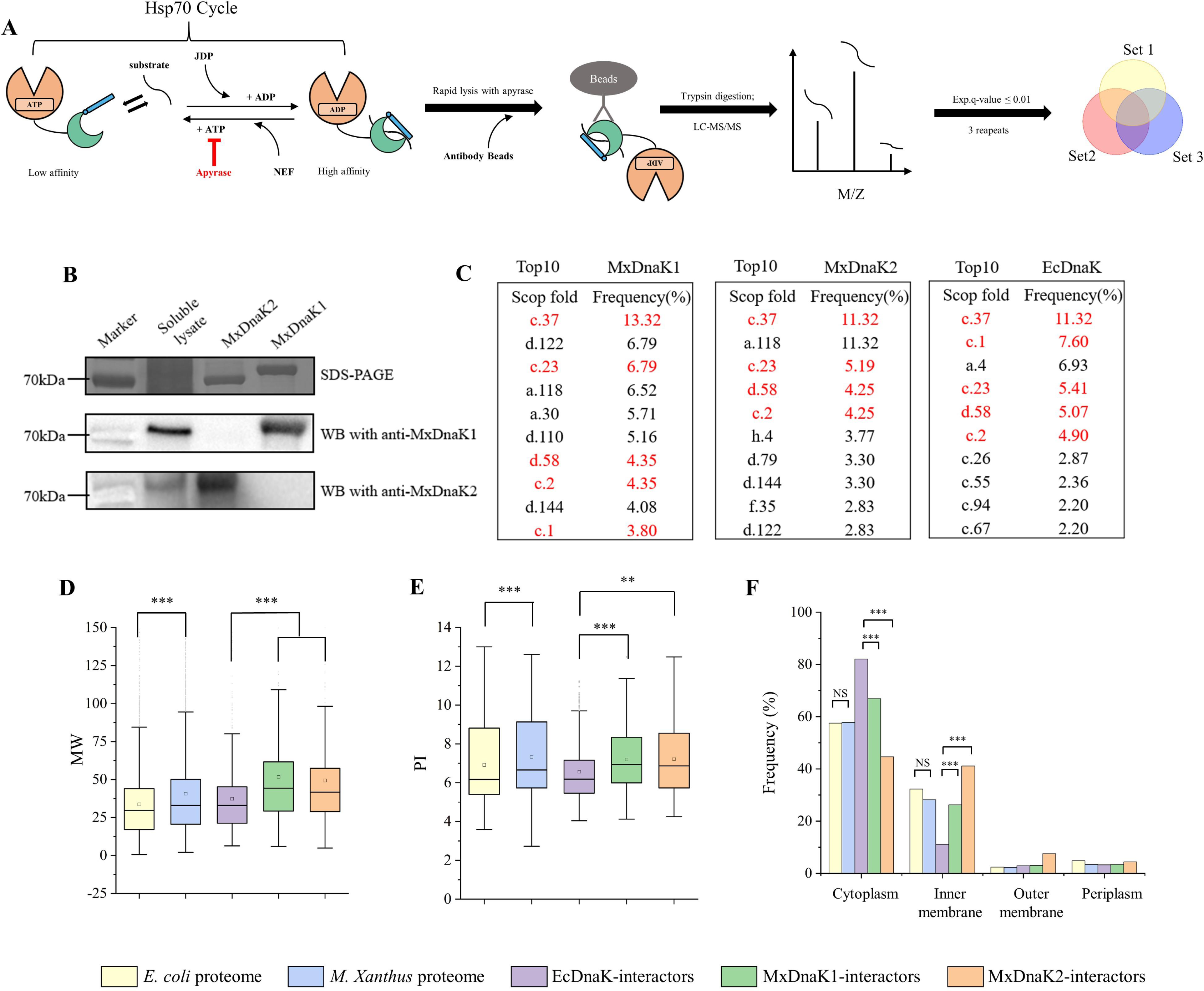
Identification and characterization of MxDnaK interactors. (**A**) A schematic diagram of Co-IP workflow for analyzing MxDnaK interactors. (**B**) Western blot detection of the antibodies with purified MxDnaK1 and MxDnaK2 proteins. The MxDnaK2 blotting band was weaker than that of MxDnaK1, consistent with their expression levels. (**C**) The top ten SCOP folds of MxDnaK and EcDnaK interactors. (**D**, **E** and **F**) Molecular weight (**D**), isoelectric point (**E**) and subcellular location (**F**) distribution of MxDnaK and EcDnaK interactors. The proteomes of *E. coli* and *M. xanthus* DK1622 served as controls. The p value in (**D**) and (**E**) is based on the Mann‒ Whitney test, and the p value in (**F**) is based on a χ^2^ test. NS: no significance, ** p < 0.01 and *** p < 0.001.

We compared the SCOP folds of the MxDnaK1 and MxDnaK2 interactors with the 674 EcDnaK interactors ^20^. The MxDnaKs and EcDnaK all preferred the α&β domain proteins, but had a low preference for the proteins containing single secondary structure units (mainly α-helices or mainly β-sheets) (**Figure S6**). For example, the proteins with SCOP fold c.37 (P-loop containing nucleoside triphosphate hydrolase) ^20^ are characterized by a complex α/β topology, and the c.37 is the top protein structure interacting with DnaK in either *E. coli* or *M. xanthus* cells. Among the top 10 preferred SCOP folds of EcDnaK (**Figure 2C**), 5 and 4 occurred in the MxDnaK1 and MxDnaK2 interactors, respectively. Similarly, of the top 10 preferred SCOP folds of MxDnaK1 and MxDnaK2, 7 were the same. Detailed comparison of the SCOP folds of the MxDnaK1 and MxDnaK2 interactors is provided in **Supplementary dataset 3**. Thus, the DnaK interactors had a similar structure classification, not only between the MxDnaKs and EcDnaK, but also between the MxDnaK1 and MxDnaK2, probably due to the high conservation of DnaK proteins ^29^.

Notably, compared to that of EcDnaK, the interactors of MxDnaK1 and MxDnaK2 were both shifted to large sizes significantly (**Figure 2D**; p<0.001) and included some very large proteins (**Supplementary dataset 3**). For example, the proteins larger than 100 kDa occupied only ∼2.7% of the EcDnaK interactors, but respectively reached ∼7.8% and ∼7.2% of the MxDnaK1 and MxDnaK2 interactors. However, there was no significant difference in the protein size interacted by MxDnaK1 and MxDnaK2 (p = 0.287). The interactor size shift was consistent with the significant increase of the molecular weights (MW) of *M. xanthus* proteome (**Figure 2D**, p<0.001); the proportions of large proteins (>100 kDa) in the *M. xanthus* and *E. coli* proteomes were 4.2% and 1.9%, respectively. Similarly, the isoelectric point (pI) values of *M. xanthus* DK1622 proteome and the MxDnaK interactors were both significantly higher than that of *E. coli* MG1655 proteome and the EcDnaK interactors (**Figure 2E**).

Although the proteomes of *M. xanthus* and *E. coli* showed similar subcellular location distributions, the interactors of MxDnaKs and EcDnaK were distributed differently (**Figure 2F**). Approximately 80% of the EcDnaK interactors were predicted to be cytosolic, while ∼11% were inner membrane proteins and ∼3% were located at the outer membrane. For the MxDnaK1 and MxDnaK2 interactors, the proportion of cytoplasmic proteins decreased to ∼65% and ∼45%, with the increase of the cytomembrane proteins to ∼25% and ∼40%, respectively. These results indicated that MxDnaKs, especially MxDnaK2, have evolved to preferentially interact with membrane proteins over EcDnaK.

### Substrate composition of the MxDnaK1 and MxDnaK2 interactomes

Of the substrate clients interacted by MxDnaK1 and MxDnaK2 (not including the JPD and NEF cochaperones), 118 were bound to both MxDnaK1 and MxDnaK2 (MxDnaK1/2 substrates), while 420 and 251 were bound exclusively to MxDnaK1 and MxDnaK2 (MxDnaK1-specific substrates and MxDnaK2-specific substrates), respectively (**Figure 3A**). These three substrate groups displayed similar M_W_ and pI distributions (**Figure S7**). However, the MxDnaK1- and MxDnaK2-specific substrates were significantly different with respect to the grand average of hydropathy (Gravy) (**Figure 3B**, P<0.001). Overall, the MxDnaK1-specific substrates were less hydrophobic than the MxDnaK2-specific substrates, which indicated that MxDnaK1 preferred to interact with thermo-sensitive proteins than MxDnaK2 ^30, 31^. This result seems to be consistent with that MxDnaK1 is induced while MxDnaK2 is reduced by heat shock ^10^.

**Figure 3.**
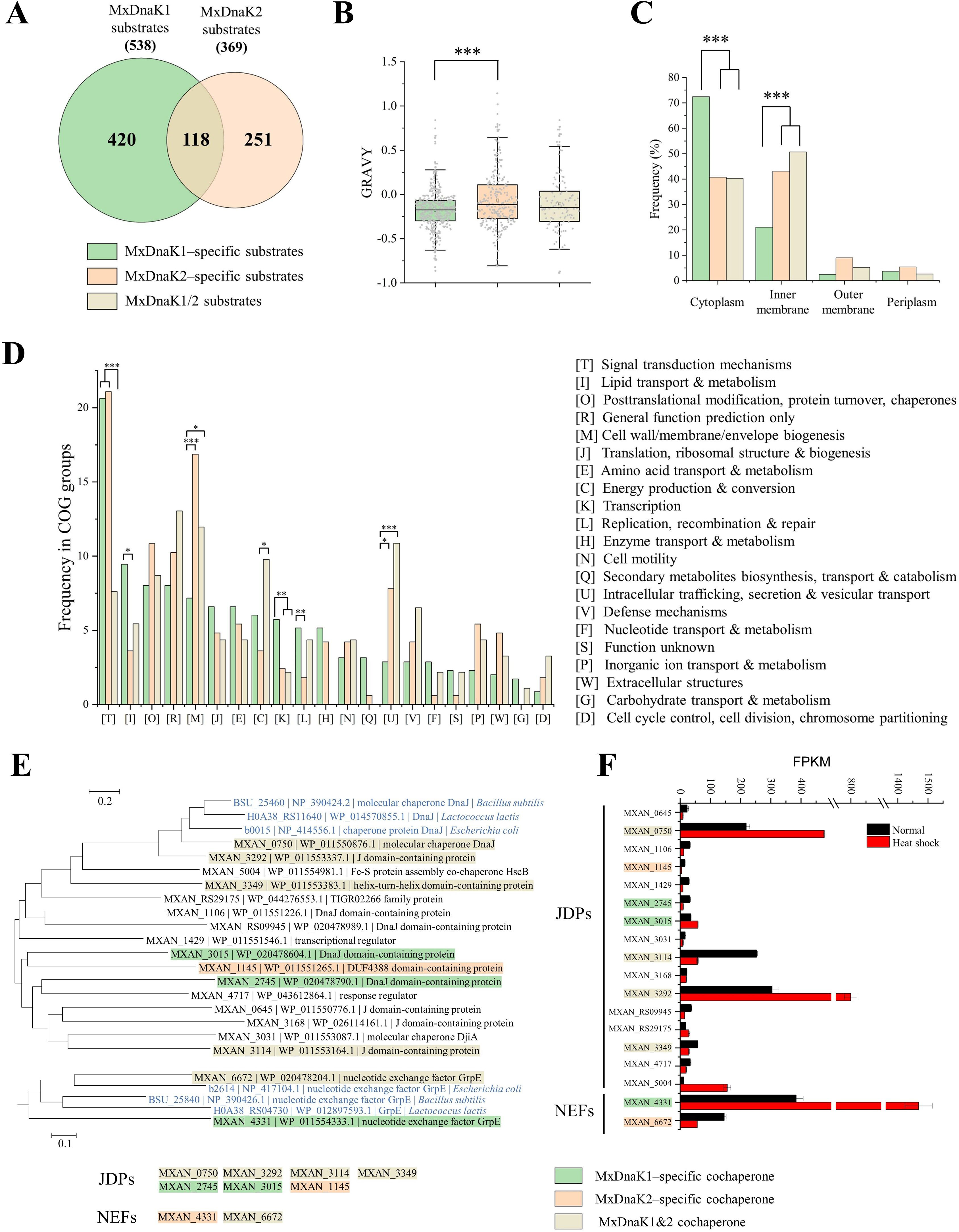
Analysis of MxDnaK1 and MxDnaK2 substrates. (**A**) Venn diagram of substrates of MxDnaK1 and MxDnaK2. (**B** and **C**) Average hydrophobicity (**B**) and subcellular localization (**C**) of MxDnaK1–specific, MxDnaK2–specific and MxDnaK1/2 substrates. (**D**) COG of protein categories of MxDnaK substrates. The p value in (B) is based on a χ2 test, in (C) and (D) is based on the Mann‒Whitney test. *p < 0.05, **p < 0.01 and ***p < 0.001. (**E**) Phylogenetic tree of the JDPs and NEFs in DK1622. The typical JDPs and NEFs from *E. coli*, *B. subtilis* and *L. lactis* (marked in blue) were employed for comparison. (**F**) Transcription levels of the DnaK cochaperone genes in *M. xanthus* DK1622 under normal growth and heat shock conditions. FPKM represents the number of fragments per kilobase of transcript sequence per million base pairs sequenced.

Another interesting difference lies in the subcellular location of substrates. We previously determined that MxDnaK2 involves in the oxidative stress of *M. xanthus* DK1622 cells ^10^. As expected, MxDnaK2 preferred to interact with membrane proteins more than MxDnaK1 (P<0.001), while the subcellular location distribution of MxDnaK1/2 substrates was similar to that of MxDnaK2-specific substrates (**Figure 3C**). Comparatively, ∼72% of the MxDnaK1-specific substrates were cytosolic proteins, and ∼20% were inner membrane proteins; this is similar to the proportion distribution of EcDnaK substrates ^20^. The proteome of *M. xanthus* is much larger than that of *E. coli*; *M. xanthus* has two duplicated DnaKs, and *E. coli* has one. Thus, we suggested that, to efficiently work with the increased proteins, MxDnaK2 has evolved more specifically to interact with membrane proteins, while MxDnaK1 persists more in the classical functions.

We predicted the substrate functions with the database of Clusters of Orthologous Genes (COGs), and the prediction ratios were 0.82 and 0.65 for the MxDnaK1 and MxDnaK2 substrates, respectively, suggesting that MxDnaK2 interacted with more function-unknown proteins than MxDnaK1. The MxDnaKs both played important roles in signal transduction of *M. xanthus* by having the highest proportion of the signal transduction substrates (COG class T in **Figure 3D**). Comparatively, the MxDnaK1-specific substrates were preferentially involved in cellular processes of lipid transport & metabolism (COG class I) and transcription (COG class K), while the MxDnaK2-specific and MxDnaK1/2 substrates both preferred the membrane-associated proteins, including cell wall/membrane/envelope biogenesis (COG class M) and intracellular trafficking, secretion & vesicular transport (COG class U). We also analyzed the Gene Ontology (GO), which showed that the MxDnaK1 clients were enriched in the Molecular Function (MF), like the proteins for adenyl nucleotide binding, ATP binding and adenyl ribonucleotide binding, while the MxDnaK2 clients were majorly located in the Biological Process (BP) group, involving transport and location (**Figure S8**). Thus, MxDnaK1 and MxDnaK2 have evolved to play divergent roles in different cellular processes, which seems to be consistent with the deletion phenotypes, i.e. the MxDnaK1 deletion is lethal, while the deletion of MxDnaK2 led to cellular sensitivity to oxidative stress and deficiency in social behaviors ^10^.

In *M. xanthus* DK1622, a total of 16 proteins were predicted to be JDPs, and none of them are neighbored by either the MxDnaK1 or MxDnaK2 gene. Seven JDPs were identified in the Co-IP assays: four (MXAN_0750, MXAN_3292, MXAN_3114 and MXAN_3349) were observed to bind to both MxDnaK1 and MxDnaK2, while two (MXAN_2747 and MXAN_3015) and one (MXAN_1145) bound exclusively to MxDnaK1 and MxDnaK2, respectively. The interactions between MxDnaK and each of the seven JDPs were further confirmed by the split NanoBit luciferase (nLuc) assay ^32^ (**Figure S9)**. Thus, MxDnaK1 and MxDnaK2 were each able to interact with several JDPs, and some JDPs were shared by the two chaperones. A phylogenetic analysis of the 16 JDPs based on their J domain sequences showed that two shared JDPs (MXAN_0750 and MXAN_3292) were clustered in the same subbranch with the typical JDP proteins (class A JDPs) of *E. coli*, *Bacillus subtilis*, and *Lactococcus lactis* (**Figure 3E**). The genes for these two JDPs exhibited high transcriptional levels and positive heat-shock responses (**Figure 3F**), which is consistent with that of MxDnaK1. However, the other two shared JDPs (MXAN_3114 and MXAN_3349) decreased their expression in response to heat shock, which is similar to that of MxDnaK2. While one of the main functions of JDPs is to interact with unfolded substrates and facilitate their delivery to Hsp70^33^, some substrates identified might bind to JDPs, rather than directly to Hsp70 ^6, 8^.

*M. xanthus* DK1622 possesses only two NEFs: MXAN_4331 and MXAN_6672. Of these two cochaperones, the transcription of MXAN_4331 was highly induced by heat shock (**Figure 3F**), which is consistent with that of the classical NEFs; whereas MXAN_6672 was deduced by heat shock, which is similar to MxDnaK2. *MXAN_6672* lies upstream of *mxdnaK*2 (*MXAN_6671*), forming a complete bicistronic operon, as verified by RT‒PCR (**Figure S10**). As expected, MXAN_6672 was identified to interact with MxDnaK2. However, both NEFs were able to bind to MxDnaK1.

### Region inter-swapping between MxDnaKs and phenotype effects

To test the roles of different regions in determining functional specificity, we performed region inter-swapping between MxDnaK1 and MxDnaK2. The two *Myxococcus* DnaKs possess an almost identical linker sequence between NBD and SBDβ (refer to **Figure S3**), which permits a stable conformation of the chimeras after domainb exchange. We obtained *M. xanthus* mutants containing wide type MxDnaK1 and a chimeric MxDnaK2 swapped with the NBD, SBDβ, SBDα or CTT domain of MxDnaK1, or wide type MxDnaK2 and a chimeric MxDnaK1 swapped with the SBDβ, SBDα or CTT, but not NBD domain of MxDnaK2 (**Figure 4A**). In *M. xanthus* DK1622, MxDnaK1 is essential for cell survival, and the in-situ deletion of the gene was obtained only after an insertion of a second copy of *mxdnaK*1 in the genome at the *attB* site^10^. To verify whether the NBD region is required for the essentiality of MxDnaK1, we performed the region swapping of the in situ MxDnaK1 gene in the att::mxdnaK1 mutant (a DK1622 mutant containing a second copy of mxdnaK1), and successfully obtained the MxDnaK1 mutant swapped with the MxDnaK2 NBD region. This result suggested that the NBD region of MxDnaK1 determine the essentiality of MxDnaK1 for cell survival, which was swapped only after an insertion of a second copy of the gene in the genome at the *attB* site ^10^. Notably, because the CTT domain is missing in MxDnaK2, we generated the corresponding mutants by directly deleting CTT from MxDnaK1 (YL2205) or inserting the MxDnaK1 CTT into MxDnaK2 (YL2215).

**Figure 4.**
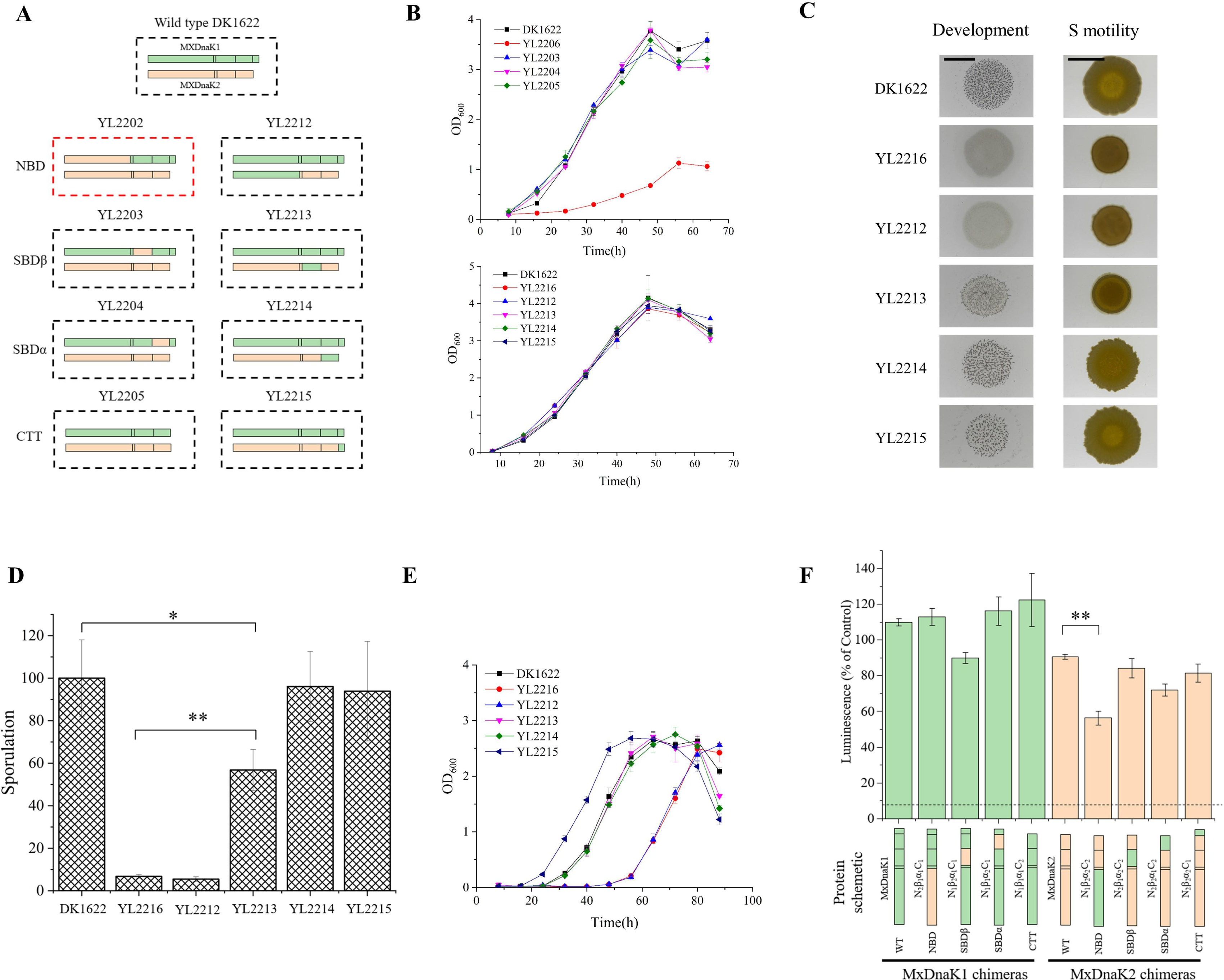
Region inter-swapping of MxDnaK1 and MxDnaK2 and their effects on cellular functions. (**A**) A schematic diagram of region inter-swapping. The swapping in YL2202 is lethal and the mutant is unavailable. (**B**) Growth curves of MxDnaK swapping mutants compared to wild-type DK1622, MxDnaK1 (YL2206) and MxDnaK2 (YL2216) deletion mutants. (**C**) Fruiting body formation (Scale bar, 4 mm) and social motility (on 0.3% agar; Scale bar, 5 mm). (**D**) Starvation-induced sporulation ability comparison (percentage to that of DK1622). (**E**) Growth curves after 1.5 h treatment with H_2_O_2_. (**F**) In-vitro holdase activities of wild-type MxDnaKs and their chimeras. The chimeras were named by N_x_β_x_α_x_C_x_. N: NBD. Β: SBDβ. α: SBDα. C: CTT. X indicates the region come from MxDnaK1 or 2. The dotted line represents the luminescence of the negative control, in which luciferase was heat treated with no chaperone. The p value in (**D**) and (**F**) is based on two tail t-test. *p < 0.05, **p < 0.01.

We analyzed the growth abilities of these region swapping mutants by comparing them with the wild-type strain, the *mxdnaK2* deletion mutant and the *mxdnaK1* depletion mutant as controls. The *mxdnaK1* depletion mutant (YL2206) was constructed by replacing the promoter of MxDnaK1 in DK1622 with a weak promoter J23112, which resulted in only ∼5% expression of MxDnaK1 (**Figure S11**). As shown in **Figure 3B**, YL2206 exhibited a significant growth defect, whereas all the inter-swapping mutants of MxDnaKs, as well as the *mxdnaK2* deletion mutant, had an almost identical growth curve to that of DK1622. These results indicated that the high expression of MxDnaK1 is required for the normal growth of *M. xanthus* cells, while the inter-swapping mutants of MxDnaKs, if viable, have no effects on cellular growth.

We previously determined that the deletion of *mxdnaK*2 (strain YL2216) affected sporulation (∼5% of wild-type), fruiting body formation, S-motility, as well as the response to oxidative stress ^10^. Among the MxDnaK2 region swapping mutants, the chimera with the NBD of MxDnaK1 (strain YL2212) showed almost identical phenotypes as the *mxdnaK*2 deletion mutant (**Figures 4C-4E**), which indicated that the NBD of MxDnaK2 was irreplaceable for the cellular functions that MxDnaK2 involved. The MxDnaK2 swapping strain with the MxDnaK1 SBDβ (strain YL2213) also defected in the phenotypes of S-motility, sporulation ability (weaker than that of the NBD swapping strain YL2212), but retained the ability to form fruiting bodies and oxidation resistance. These results indicated that the SBDβ swapping significantly, although not completely, limited the cellular functions of MxDnaK2. Comparatively, the exchange of SBDα or CTT had no obvious phenotypic impacts.

To determine whether the MxDnaK chimeras hold the chaperone functions, we assayed the *in vitro* holdase activity in protecting luciferase proteins from aggregation, which is often employed to measure the cochaperone-independent chaperone activities of DnaK ^35^. We expressed and purified the MxDnaK1 and MxDnaK2 proteins and their mutants. The MxDnaK1 and MxDnaK2, despite having divergent cellular functions *in vivo*, exhibited similar *in vitro* holdase activities at various concentrations (**Figure S12**). Interestingly, all the variants, including N_2_β_1_α_1_C_1_ (the MxDnaK1 chimera with the MxDnaK2 NBD), showed similar holdase activity to the wild-type protein except for N_1_β_2_α_2_C_2_ (the MxDnaK2 chimera with the MxDnaK1 NBD), which produced a relatively weak luminescence of heat-denatured luciferase (**Figure 3F**). Thus, the region inter-swapping chimeras of MxDnaK1 or MxDnaK2 still retained chaperone activities, at least the holdase ability.

### Effects of NBD and SBDβ on the interactomes of MxDnaK1 and MxDnaK2

To explore possible contributions of NBD and SBDβ to the substrate recognition, we performed Co-IP with N_1_β_2_α_1_C_1_, N_1_β_2_α_2_C_2_ and N_2_β_1_α_2_C_2_ in YL2203, YL2212 and YL2213 to identify their substrates, respectively (**Supplementary dataset 3**). Surprisingly, in comparison with the 538 and 369 substrates of wild-type MxDnaK1 and MxDnaK2, the substrate numbers of N_1_β_2_α_1_C_1_, N_1_β_2_α_2_C_2_ and N_2_β_1_α_2_C_2_ decreased to 150, 54 and 141, respectively. The results suggested that the swapping of NBD or SBDβ, especially the former, greatly decreased the interaction between substrates and DnaKs, although the cells with chimeric DnaKs exhibited no effect on growth.

Although the number decreased sharply, the substrates of these three chimeras overlapped extensively with that of the wild-type MxDnaKs: ∼70%-82% substrates of the chimeras were in the substrate pools of MxDnaK1 or MxDnaK2 (**Figure 5A**). The N_1_β_2_α_1_C_1_ substrates consisted of not only some of the original MxDnaK1 substrates but also some MxDnaK2-specific substrates. Similar results were obtained with N_1_β_2_α_2_C_2_ and N_2_β_1_α_2_C_2_ substrates. Notably, in contrast to the 22%-32% overlapping ratio of the substrates between MxDnaK1 and MxDnaK2, 74 proteins were overlapped in the interactomes of N_1_β_2_α_1_C_1_ and N_2_β_1_α_2_C_2_, accounting for ∼50% of their respective total substrates (refer to **Supplementary dataset 3**). Thus, the reduced substrates by the chimeras were still those essential components that are required for cellular growth, and the lost were mainly related to some specific cellular functions like sporulation. In addition, some new substrates that were absent in the MxDnaK1 or MxDnaK2 interactomes were recognized by each of the three chimeras. These results not only validated the contribution of NBD and SBDβ in substrate recognition but also suggested that the swapping endowed the chimeras with some new functions.

**Figure 5.**
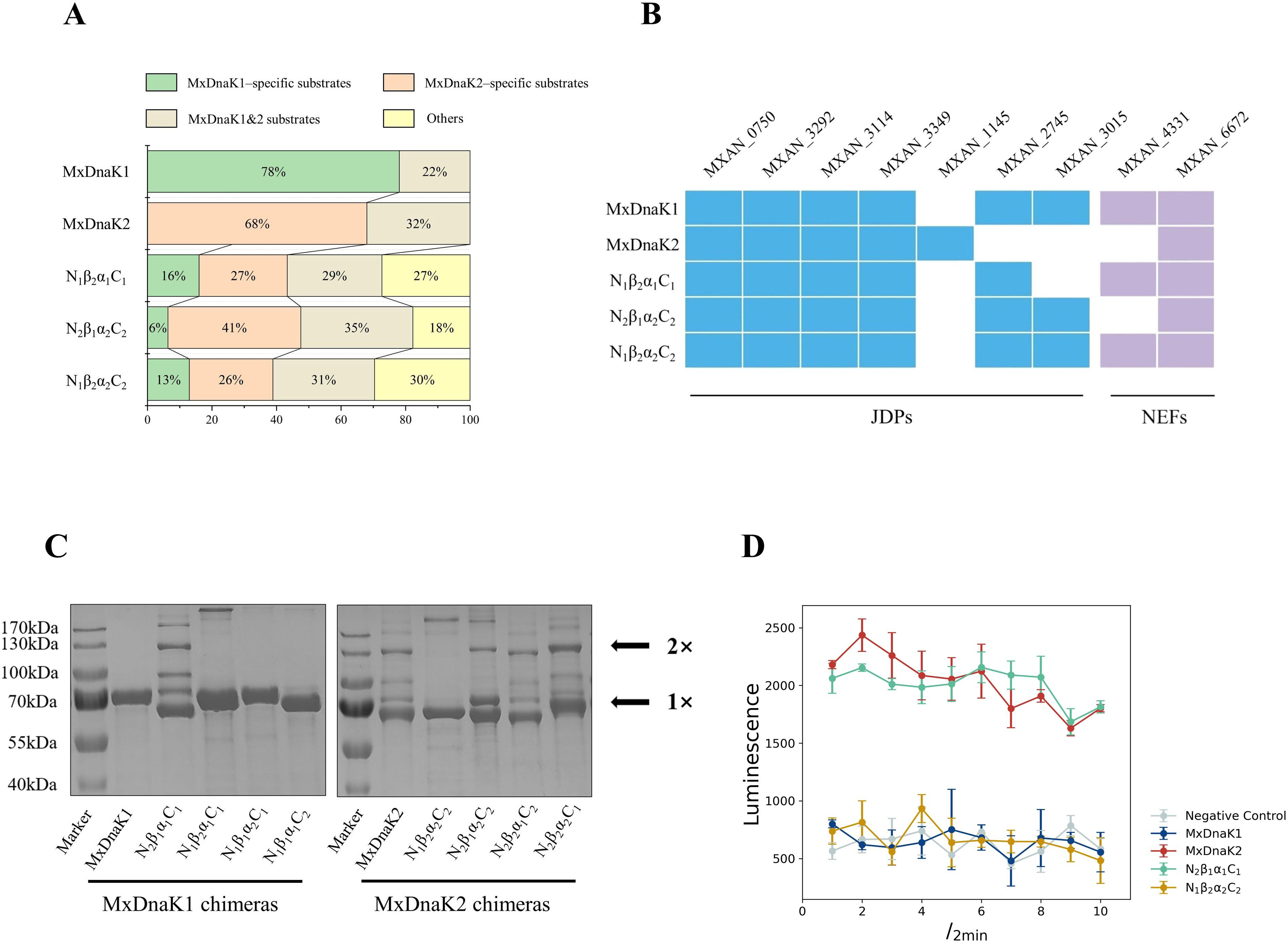
Effects of exchanging NBD and SBDβ. (**A**) Class distribution of chimera substrates compared to that of wild-type MxDnaK substrates. (**B**) Cochaperones distribution interacted by chimeras. (**C**) SDS‒PAGE analysis of wild-type MxDnaKs and their chimeras, showing their oligomeric states. The positions of the monomer (1×) and dimer (2×) are indicated by arrows. The two distinct bands observed in the purified His6-MxDnaK2 lane, falling within the molecular weight range of 70-100 kDa, were identified as the interaction between His6-MxDnaK2 (65.3kDa) and anti-FlhDC factor (WP_001300634.1, 27 kDa) as well as the 50S ribosomal protein L28 (WP_000091955.1, 9 kDa). (**D**) Split NanoLuc luciferase analysis of MxDnaKs and their NBD swapping chimeras. Negative controls were performed with empty vectors.

In terms of cochaperones, the three chimeras were found to interact with all the four MxDnaK1/2 JDPs (MXAN_0750, MXAN_3292, MXAN_3114 and MXAN_3349) but varied with the MxDnaK1/2-specific JDPs (**Figure 5B**). For example, the NBD- and SBDβ-swapped MxDnaK2 chimeras (N_1_β_2_α_2_C_2_ and N_2_β_1_α_2_C_2_) were both unable to recognize the MxDnaK2-specific JDP (MXAN_1145), but bound to MxDnaK1-specific JDPs (MXAN_2745 and MXAN_3015). In N_1_β_2_α_1_C_1_, the exchange of SBDβ abrogated the binding to MXAN_3015, but still accepted MXAN_2745. In comparison, the exchange of SBDβ did not change the recognition of MxDnaK1 or MxDnaK2 to NEFs, while N_1_β_2_α_2_C_2_ bound to the same NEFs (both MXAN_4331 and MXAN_6672) as MxDnaK1. These results suggested that NBD plays an important role in NEF or JDP recognition, while SBDβ affected the recognition of JDPs only; the recognition of NEF and JDP did not affect each other.

In *E. coli*, the DnaK proteins were able to form a small amount of dimer via the disulfide bonds, which could be reduced by DTT ^35, 36^. Interestingly, in the experiments of expression and purification of the MxDnaK proteins, we found that MxDnaK2 formed an obvious dimer band, while MxDnaK1 presented only as a monomer band (**Figure S13** and **Figure 5C**). We treated the MxDnaK2 proteins with 100 mM DTT, and the MxDnaK2 dimer was dissociated to the monomer form, and the band compositions were confirmed by LC/MS. We further tested the oligomeric states of MxDnaK1 and MxDnaK2 *in vivo* by using the nLuc assays ^32^. The MxDnaK proteins were fused with either large (LgBiT) or small (SmBiT) fragment of nLuc, and the two kinds of fused proteins were co-expressed in an *E. coli* host. The formation of dimers was monitored by the reconstituted nLuc activity. Consistent with the *in vitro* results, MxDnaK2-nLuc exhibited obvious nLuc activity, while MxDnaK1-nLuc showed no difference from the negative control (**Figure 5D**). The above results indicated that MxDnaK2, but not MxDnaK1, is prone to form the dimer form either *in vitro* or *in vivo*. Interestingly, the MxDnaK2 chimera with the NBD region of MxDnaK1 (N_1_β_2_α_2_C_2_) could not form the dimer band, while the MxDnaK1 chimera with the NBD region of MxDnaK2 (N_2_β_1_α_1_C_1_) formed an obvious dimer band (**Figure 5C**). The *in vivo* nLuc assays also confirmed the dimeric state changes (**Figure 5D**). Thus, the NBD region is fully responsible for the difference of the dimeric state of MxDnaK1 and MxDnaK2, which is probably correlated with their substrate interacting activities, as well as the cochaperones.

## Discussion

The cellular proteomes of eukaryotes are more complex than those of prokaryotes, and the proteomic complexities in prokaryotes are also in hierarchy. Hsp70 proteins are among the most ubiquitous chaperones that play important roles in maintaining proteostasis and resisting environmental stresses. To meet the increased protein-refolding requirement, cells across all eukaryotes and ∼40% prokaryotes encode multiple Hsp70 family members ^8^. Although might be functionally redundant, Hsp70 family members generally exhibit a high degree of specialization. The differential expression, tissue-specific abundance and subcellular localization of distinct Hsp70 isoforms are all suggested to contribute to diversification of the cellular functions of Hsp70s in eukaryotes ^37, 38^. We found that the duplication of canonical Hsp70s (DnaKs) is positively correlated with the increase of proteomic complexity, and the bacteria that have evolved more and larger proteins for their complex lifecycles, such as myxobacteria, are prone to encode duplicate DnaK proteins, helping cells maintain proteome integrity.

Due to high conservation, different Hsp70 paralogs are often regarded functionally interchangeable ^11, 39^. However, functional specificity does exist between Hsp70 paralogs, consequently playing different cellular roles, e.g., the cytosolic SSAs and SSBs in yeast^16^, the DnaK, HscA and HscC in *E. coli* ^40^, as well as the MxDnaK1 and MxDnaK2 in *M. xanthus* ^10^. Although the two DnaKs of *M. xanthus* DK1622 both retain some fundamental functions, they are functionally different, which is consistent with their largely nonoverlapping substrates. We found that the substrates interacted by MxDnaK1 and MxDnaK2, or MxDnaKs and EcDnaK, have similar structures, molecular weights and isoelectric points. MxDnaK1 prefers to bind to proteins without an extensive hydrophobic core, which are more unstable and aggregation prone in response to heat shock ^31^. In comparison, the MxDnaK1/2 substrates and MxDnaK2-specific substrates prefer membrane proteins, which is consistent with their COG or GO functional assignment. The subcellular location distribution of MxDnaK1-specific substrates was more like that of EcDnaK substrates. Furthermore, typical DnaK proteins are usually characterized by a high expression level and a positive heat shock response. For the two MxDnaKs in *M. xanthus*, MxDnaK1 expressed greatly higher than MxDnaK2 (approximately eight times); the transcription of MxDnaK1 was increased, while that of MxDnaK2 was decreased by heat shock ^10^. Thus, MxDnaK1 exhibits similar transcriptional regulation and substrate preference as canonical DnaKs, while MxDnaK2 has evolved for some unique cellular processes of myxobacteria.

Region swapping was previously performed to explore the effects of Hsp70s on phenotypes and specific substrates in yeast or human cells ^41, 42^. In this study, we performed region swapping between MxDnaK1 and MxDnaK2, and assayed their effects on cellular phenotypes. In addition, we checked the substrate pools of MxDnaK1 and MxDnaK2 chimeras to clarify the relationships between the structural and functional divergence. Of the Hsp70 regions, NBD and SBDβ are highly conserved in evolution, and are also important for binding substrates; whereas SBDα and CTT are not only rather variable, but also interchangeable with small effects on the chaperone functions. Due to the highly conserved working mechanism and different peptide-binding sites, it has been previously hypothesized that SBDβ plays a central role in the functional specificity of Hsp70 paralogs ^43, 44^. The importance of NBD as a driver of functional specificity has also been experientially proven in human and yeast Hsp70s ^41, 45^. In agreement with these findings, we found that SBDβ was interchangeable between MxDnaK1 and MxDnaK2, while the exchange of NBD led to similar phenotypes as the deletion of MxDnaK1 or MxDnaK2. Nevertheless, the NBD-swapped MxDnaK chimeras (N_1_β_2_α_2_C_2_ and N_2_β_1_α_1_C_1_) still retained their holdase function *in vitro*, and the region swapping chimeras bound similar substrates, although the substrate numbers were greatly reduced. These results indicated that the exchange of NBD, as well as SBDβ, did not completely denature MxDnaK1 or MxDnaK2.

Perhaps one of the most surprising findings is the different oligomeric states of MxDnaKs. The multitude of functions performed by Hsp70s, in addition to the Hsp70-outside cooperation with other cooperating protein folding machineries like Hsp60s and Hsp90s, are specified by the Hsp70-inside cooperation with different family members, as well as the JDP and NEF cochaperones. The JDP and NEF cochaperones changed in the NBD swapping chimeras, but only JDP cochaperone changed in the SBDβ swapping chimeras; these results are consistent with the interactions of these two domains, i.e. JDP binds at the interface between the NBD and the SBDβ while NEF binds specifically to the NBD region ^8^. The corresponding changes in the substrate spectrums were thus suggested to be related not only to the shifts of cochaperones, but also to the changes in domains. MxDnaK2, not MxDnaK1, is prone to form dimer both *in vitro* and *in vivo*. The Hsp70 oligomer has been confirmed to have a limited foldase activity, and maintain the holdase activity ^46^. In addition, mutation of the Hsp70 dimerization interface significantly decreased the Hsp70-JDP affinity in both bacteria and mammals ^36, 47^. It is unsurprising that Hsp70 dimerization impedes the association with cochaperones and accordingly leads to defects in co-chaperone associated refolding activity ^48^. Similarly, our results demonstrated that exchange of the NBD not only reversed the oligomeric states of MxDnaK1 and MxDnaK2, but also exchanged their individual JDPs. JDPs play a crucial role in this process as they possess the ability to bind substrates and facilitate their targeted delivery. Hence, the oligomeric states, which consequently impact DnaK interactions with JDPs, also contribute to the disparities in substrate recognition between MxDnaK1 and MxDnaK2.

In this study, for the first time, we characterized the interactomes of bacteria Hsp70 paralogs, and determined the importance of NBD and SBDβ in their functional diversity based on phenotype and substrate identification. The reasons for the importance and cooperation of different regions are still unclear, of which our studies provide some important clues. The multitude of functions performed by Hsp70s are specified through multilayered networks of JDP and NEF cochaperones. Indeed, cochaperones have been reported to influence the functional diversity of Hsp70 in eukaryotes ^33, 38^. For example, the diverse preference of NEFs leads to opposing effects of HSPA1A and HSPA1L on the fate of superoxide dismutase 1 (SOD1) mutants ^42^. We also found that NBD and SBDβ played important roles in JDP and NEF recognition of MxDnaK. Thus, it will be interesting to further explore the relationships between the cochaperone recognition and the structural and functional divergence of Hsp70s.

## Methods and materials

### Distribution analysis of *dnaK*s in prokaryotes

The 16083 representative prokaryotic genomes were obtained from the NCBI Reference Sequence Database. Conserved domain information of DnaKs and taxonomic information of prokaryotes were obtained from the Conserved domain database ^49^ and NCBI taxonomy database. All DnaK proteins were identified using RPS-BLAST based on the conserved domain (PRK00290), and the retrieval condition was set to an E value < 0.01. The amino acid sequences were aligned using the protein sequence alignment program in MAFFT ^50^, and the *dnaK* gene sequences from representative prokaryotic genomes were retrieved from the NCBI database. The Ka/Ks values among the *dnaK* duplicate pairs were calculated using KaKs_Calculator 3.0 ^51^ with the MLWL ^52^. Phylogenetic trees were constructed and annotated by MEGA, CVtree3 and iTOL online.

### Strains, plasmids, and growth conditions

The strains, plasmids and primers used in this study are listed in **Tables S2 and S3**. The *M. xanthus* strains were cultivated in CTT medium at 30°C. The *E. coli* strains were routinely grown in Luria-Bertani (LB) medium at 37°C. When required, 100 μg/ml ampicillin, 40 μg/ml kanamycin, 10 μg/ml tetracycline or 20 μg/ml chloromycetin (final concentrations; Solarbio) was added to the solid or liquid medium.

### Hsp70 Protein purification

Genomic DNA from *M. xanthus* DK1622 and region swapping mutants served as a template for the PCR amplification of the *hsp70* genes. Subsequently, the genes were cloned into the pET28A vector and were fused with an N-terminal His tag gene. These constructs were then overexpressed in Escherichia coli BL21 (DE3) cells upon induction with 0.1 mM isopropyl β-D-1-thiogalactopyranoside (IPTG). After incubation at 16°C for 22 h, the BL21 cells were resuspended in resuspension buffer (25 mM Tris-HCl, 200 mM NaCl, 5% (v/v) glycerol; pH 8.0) and sonicated on an ice-water slurry. The cell lysate was centrifuged at 12000 rpm for 30 min to remove the debris, and then the supernatant was incubated with Ni^2+^-NTA (GE Healthcare) at 4°C for 2 h. After incubation, the Ni Sepharose was washed with resuspension buffer supplemented with 20 mM imidazole to remove nonspecific binding proteins, and the Hsp70 proteins were eluted with elution buffer (25 mM Tris-HCl, 200 mM NaCl, 5% (v/v) glycerol, 250 mM imidazole, pH 8.0). If necessary, centrifugal concentrators (Millipore) were used to concentrate purified proteins. And thrombin was used to removed His_6_ tag.

### Western blot and antibodies

WB was performed as described with minor modifications ^53^. Briefly, proteins were loaded into 12% SDS‒PAGE gels and then transferred onto activated PVDF membranes. The PVDF membranes were incubated with anti-Hsp70 antibodies in TBST buffer (20 mM Tris-HCl, 500 mM NaCl, 0.05% (v/v) Tween 20; pH 7.5). Then, the membranes were washed six times with TBST buffer and blotted with horseradish peroxidase (HRP)-conjugated goat anti-rabbit secondary antibodies (Sigma). After washing with TBST buffer again, visualization was performed with enhanced chemiluminescence (ECL) detection reagents (GE Healthcare) and a ChemiDoc Imaging System (Bio-Rad) with Image Lab (Bio-Rad) software.

Anti-Hsp70 antibodies were prepared by ABclonal Biotechnology Co., Ltd, China.

### Isolation of Hsp70-Interactor Complexes

*M. xanthus* strains were grown for 24 h in CTT medium to the midlogarithmic phase (∼1 OD_600nm_). Then, the cultures were cooled and harvested by centrifugation at 8000 rpm for 5 min. The cell pellets were washed twice with PBS and suspended in lysis buffer (10 mM Tris-HCl, 0.1% (v/v) Triton X-100, 10 mM MgCl_2_, 12.5 U/ml benzonase, 1x EDTA-free protease inhibitor cocktail, 50 U/ml apyrase; pH 8.0) ^27^. After sonication on ice, the debris was removed by centrifugation (12000 rpm, 30 min), and the supernatant was used as the starting material for coimmunoprecipitation (Co-IP). Co-IP was performed using the Protein A/G Matrix Immunoprecipitation Kit (Beaverbio, China). Briefly, the supernatant was incubated at 4°C overnight with anti-Hsp70 antibodies and beads. Then, the bead-antibody-Hsp70-Interactor complexes were washed three times with IP Washing buffer (Beaverbio), and the Hsp70-Interactor complexes were finally eluted from the beads with IP Elution buffer (Beaverbio). We performed the Co-IP experiments three times, and the proteins identified in at least two of three independent experiments were regarded as the Hsp70 interactors.

Eluates were treated with the FASP method with minor modifications. Briefly, the samples were washed two times with UA buffer (8 M urea, 0.1 M Tris pH 8.5) and then incubated at 56°C for 1 h with 100 mM DTT in UA buffer to reduce disulfide bonds. After DTT treatment, samples were alkylated with 50 mM iodoacetamide (IAA) by incubation for 30 min in the dark. Then, the samples were washed three times with UA buffer and two times with ammonium bicarbonate (ABC) buffer (50 mM NH_4_HCO_3_). After washing, trypsin (Sigma) was added at a final concentration of 0.5 μg/ml, and the samples were incubated at 37°C overnight. Peptides were eluted with ABC buffer and then desalted with C18 tips (Millipore) according to the manufacturer’s protocol. The final samples were lyophilized at room temperature and resuspended in ultrapure water for LC‒MS analysis.

### Protein identification and bioinformatic analysis

Protein identification by LC‒MS/MS was conducted on Orbitrap Fusion Lumos (Thermo Fisher) by Shandong University at core facilities for life and environmental sciences. Raw files were analyzed by Proteome Discoverer software, which comes with the instrument. The interactors were identified with an Exp.q-value ≤ 0.01. We implemented negative controls for each MxDnaK in order to eliminate potential non-specific interactions with Protein A/G beads or antibodies. Specifically, we conducted a CO-IP experiment without the presence of antibodies to evaluate any non-specific binding to the Protein A/G beads. Additionally, the mxdnak2-deleted mutant (strain YL2216) and the MxDnaK1 swapping strain with the MxDnaK2 SBDα (strain YL2204) were utilized to investigate the non-specific binding to the antibodies of MxDnaK2 and MxDnaK1, respectively. As the SBDα of MxDnaK1 was employed as antigen to generate antibodies, and YL2204 can’t be recognized by anti-MxDnaK1 (Figure S5).

Protein fold assignment was derived from the SCOPe (Structural Classification of Proteins — extended) 2.07 superfamily database (https://scop.berkeley.edu/) ^28, 54^. Sublocation of client proteins was predicted with pSORTb 3.0 (https://db.psort.org/) ^55^. Theoretical isoelectric points (pIs) and molecular weights (MWs) of protein sequences were predicted with Expasy (https://web.expasy.org/compute_pi/) ^56^. Functional assignment of client proteins was performed using the COGs database (Database of Clusters of Orthologous Genes, https://www.ncbi.nlm.nih.gov/research/cog) and GO database (Gene ontology database, http://geneontology.org/) ^57, 58^. The substrate features of EcDnaK were provided in a previous study ^20^. Proteome information of *E. coli* MG1655 and *Myxococcus xanthus* DK1622 was obtained from NCBI, and the protein features were predicted as mentioned above.

### Split NanoLuc luciferase (nLuc) oligomerization assay

NanoBit luciferase is split into two fragments: SmBiT and LgBiT ^32^. Either SmBiT or LgBiT fragment was fused to the C terminus of a MxDnaK, and the MxDnaK-SmBiT and MxDnaK-LgBiT fragments were cloned into the pET28A and pACYC Duet-1 plasmids, respectively. Then, the pET28A-MxDnaK-SmBiT and pACYC-MxDnaK-LgBiT expression vectors were transformed into the *E. coli* BL21(DE3) cells simultaneously. The negative control contained the empty pET28A and pACYC vectors without the MxDnaK fragments. The luminescence was monitored using the Nano-Glo® Live Cell Assay System kit (#N2011, Promega).

Additionally, pET28A-JDP-SmBiT and pACYC-MxDnaK-LgBiT expression vectors were used to validate the interaction between DnaKs and JDPs. And αSyn served as the positive control ^59^.

### RT‒PCR analysis

*M. xanthus* DK1622 cells were collected from 24-h cultures, which were inoculated into CTT medium at a final concentration of OD_600_ 0.04. The cultures were harvested after 24 h, and the RNA was extracted immediately using a bacterial RNA extraction kit (TaKaRa). The purified RNA extracts were reverse-transcribed to cDNA. The cotranscription of *mxdnak2* and *MXAN_6672* was tested by primers.

### Region swapping

Region swapping of *hsp70* genes in *M. xanthus* was performed using positive-negative KG cassettes ^54^. The upstream and downstream homologous arms and different domain sequences were cloned from the DK1622 genome and fused with pBJ113 to form region-swapping plasmids, which were then transferred via electroporation into *M. xanthus* DK1622 cells as previously described. The selected colonies were inoculated onto CTT agar plates supplemented with 1.5% galactose (Sigma) for a second round of screening. The region swapping mutants were identified based on their galactose resistance and kanamycin sensitivity phenotypes, as well as by PCR and sequencing verification. The CTT domain is missing in MxDnaK2. Thus, exchanging the CTT of MxDnaK2 with that of MxDnaK1 was performed by directly inserting or deleting CTT in their respective regions.

### Developmental assays

The developmental assays were performed according to the method used in a previous study ^60^. Briefly, *M. xanthus* cells were harvested at midlogarithmic phase and resuspended in TPM buffer (10 mM Tris-HCl, 8 mM MgSO_4_, 1 mM K_2_HPO_4_-KH_2_PO_4_; pH 7.6) to a final cell concentration of 5×10^9^ cells/ml. Eight-microliter aliquots of concentrated cell suspension were spotted onto TPM agar. The colonies were incubated at 30°C and monitored every 24 h under a dissection microscope. Three replicates of fruiting bodies were harvested in 0.5 ml of TPM buffer and incubated at 50°C for 2 h to kill vegetative cells. After sonication, the spore suspensions were serially diluted and plated on CTT agar. After 5 days, the sporulation rate was calculated as the number of colonies. Assays were performed with three biological replicates.

### Swarm assays

The swarm assays were performed according to the method used in a previous study^61^. Briefly, *M. xanthus* cultures were harvested at midlogarithmic phase, washed three times with TPM buffer (pH 7.6), and resuspended to a final concentration of 5×10^9^ cells/ml. Aliquots (2 μl) were dropped onto 0.3% CTT agar. After 72 h incubation, the swarming size of *M. xanthus* cells was monitored.

### Growth and oxidative resistance analysis

*M. xanthus* strains were grown in CTT medium with shaking at 200 rpm at 30°C to the midlogarithmic phase (∼1 OD_600_) as seed liquid. Then, the cells were inoculated at a final cell concentration of 0.04 OD_600_ and grown in CTT medium for 64 h with shaking at 200 rpm. The OD_600_ value was read every 8 h.

To assay growth under oxidative damage stress, the seed liquid was treated with 1.5 mM hydrogen peroxide (H_2_O_2_) for 30 min before inoculation. The culture time was accordingly extended to 88 h.

### Luciferase assays

The luciferase holdase activity of Hsp70 proteins and mutants was performed as described with minor changes ^35^. Native firefly luciferase (Promega) was diluted in holdase buffer (50 mM HEPES, 300 mM KCl, 10 mM MgSO_4_, and 20 mM DTT, pH 7.5) at a concentration of 0.032 μM. After mixing with Hsp70 at a 1:1 ratio, the luciferase was heated to 39.5°C for 8 min. Fifty microliters of the reaction mixture was transferred to a 96-well, opaque assay plate, and then 50 μl of 5% (v/v) SteadyGlo reagent (Promega) in glycine buffer (50 mM glycine, 30 mM MgSO_4_, 10 mM ATP, 4 mM DTT, pH 7.8) was added to each well. Next, the luminescence at 560 nm was measured. For the experiment, the negative control contained all but Hsp70 proteins, and the 100% control contained native luciferase without heat shock.

### Statistical analysis

Significant differences were analyzed statistically based on paired *t* tests (two tailed), chi-square tests or Mann-Whitney tests by SPSS or Scipy: *p < 0.05, **p < 0.01 and ***p < 0.001.

## Acknowledgements

This work was financially supported by the National Natural Science Foundation of China (NSFC) (Nos. 32070030 and 31670076), the Key Program of Shandong Natural Science Foundation (Nos. ZR2016QZ002), Special Investigation on Scientific and Technological Basic Resources (Nos. 2017FY100302), National Key Research and Development Programs of China (Nos. 2018YFA0900400 and 2018YFA0901704) to YZL, Natural Science Foundation of Jiangsu Province (Nos. BK20220267) to LZ and Postdoctoral Foundation of Qingdao (Nos. QDBSH20220201044) to PZ.

We would like to thank Jingyao Qu, Jing Zhu and Zhifeng Li from State Key laboratory of Microbial Technology of Shandong University for help and guidance in LC‒MS/MS.

## Supplementary materials

**Figure S1.** Duplication ratio of *dnaK* in prokaryotes at the class and order levels (top 10).

**Figure S2.** Phylogenetic analysis of myxobacterial DnaK proteins. Left: The phylogenetic tree of the 93 DnaK proteins from 48 myxobacteria. Right: The phylogenomic tree based on the 48 genome sequences.

**Figure S3.** Domain organization and identity of MxDnaK1 and MxDnaK2. EcDnaK (DnaK from *E. coli*) served as a control. NBD: nucleotide binding domain; L: linker; SBDβ: substrate binding domain β; SBDα: substrate binding domain α; CTT: C terminal tail.

**Figure S4.** Western blot detection of the antibodies of MxDnaK1 and MxDnaK2 proteins. Soluble lysate represents the cell lysate of *M. xanthus* DK1622. both purified MxDnaK1 and MxDnaK2 proteins were engineered with an N-terminal His_6_ tag.

**Figure S5.** Western blot detection of YL2204. YL2204 were employed to exclude the non-specifically bound proteins of MxDnaK1 antibodies.

**Figure S6.** Secondary structure unit distribution of MxDnaKs and EcDnaK interactors. The protein fold assignment was derived from the SCOPe (Structural Classification of Proteins-extended) 2.07 superfamily database.

**Figure S7.** Molecular weight (**A**) and isoelectric point (**B**) distribution of MxDnaK1– specific substrates, MxDnaK2–specific substrates and MxDnaK1/2 substrates.

**Figure S8.** GO enrich analysis of MxDnaK1 and MxDnaK2 substrates.

**Figure S9.** nLuc detection of the interaction between MxDnaKs and JDPs. Positive and Negative controls were performed with αSyn and empty vectors, separately.

**Figure S10.** RT‒PCR detection on the co-transcription of *mxdnaK2* (*MXAN_6671*) and *MXAN_6672*. M: marker; + : positive control using the total DNA extracted from DK1622 as the template; - : negative control with no addition of reverse transcriptase; EG: experimental group.

**Figure S11.** The transcription level of *mxdnaK1* in the *mxdnaK1* depletion mutant (YL2206). The YL2206 strain was generated by replacing the native promoter with the J23112 promoter.

**Figure S12.** Holdase ability of various concentration of wild-type MxDnaK1 and MxDnaK2. The Y-axis (luminescence) was presented as the percent of control group in which the luciferase was not treated with heat shock (Control).

**Figure S13.** SDS-PAGE analysis of MxDnaK1 and MxDnaK2 with MBP or His_6_ tag. The positions of the monomer (1X) and dimer (2X) are indicated by arrows. The His_6_ and MBP tags were removed with thrombin to corroborate that they are not interfering with the dimerization.

**Figure S14.** The complete 2-hours Split NanoLuc luciferase analysis of MxDnaKs and their NBD swapping chimeras. Negative controls were performed with empty vectors. According to the working principle of the nLuc assay, the amount of fluorescent substrate is limited. Therefore, even for proteins that form oligomers, the fluorescence value gradually decreases and reaches a plateau, similar to the negative control. This gradual decline in fluorescence is a significant indicator of protein interaction.

**Table S1.** The distribution of DnaKs in prokaryotes.

**Table S2.** Bacterial strains and plasmids used in this study.

**Table S3.** List of primers used in this study.

**Dataset 1.** The occurrence of DnaKs in representative genomes of prokaryotes.

**Dataset 2.** The Ka/Ks values of duplicate DnaK pairs.

**Dataset 3.** The identified interactors and substrates of MxDnaKs.

